# Predicting stimulus representations in the visual cortex using computational principles

**DOI:** 10.1101/687731

**Authors:** Agustin Lage-Castellanos, Federico De Martino

**Affiliations:** Department of Cognitive Neuroscience Maastricht University.; Department of NeuroInformatics, Cuban Center for Neuroscience, Cuba; Center for Magnetic Resonance Research, University of Minnesota, Minneapolis, USA

## Abstract

In this report we present a method for predicting representational dissimilarity matrices (RDM). This method was used during the MIT challenge 2019. The method consists in combining perceptual and categorical RDMs with RDMs extracted from deep neural networks.

## Introduction

Understanding the computations underlying the extraction of visual information through the hierarchical organization of brain areas is one of the main focuses in computational neuroscience. In this context, the MIT-challenge 2019 asked the participants to predict the representational structure of physiological responses to visual stimuli (in the form of representational dissimilarity matrices (RDM) (Kriegeskorte et al., 2008) measured with fMRI and MEG data (Cichy et al., 2016). The MEG dataset consisted of RDM matrices from 15 volunteers extracted in two temporal windows of the electrophysiological responses: 1) 100-120 ms (early) and 2) 165-185 ms (late), sampled in twenty time points each. The fMRI dataset consisted, for the same 15 volunteers, of the RDM in two cortical regions: 1) early visual cortex (EVC) and 2) the inferior temporal cortex (ITC). The training data, consisted of two sets of RDMs (obtained from responses to 92 and 118 images respectively). The challenge’s aim was to predict the RDMs (of the early/late MEG window and the EVC/ITC fMRI responses) of a test data set, which consisted of 78 images obtained from an independent sample of subjects. In addition to the MEG and fMRI data and the visual stimuli, the organizers provided participants with the representation of the training and test images by three different deep neural networks (DNNs - alexnet, vgg, and resnet) (Khaligh-Razavi and Kriegeskorte, 2014).

## General description of the algorithm

We constructed four initial estimations of the RDM matrices (two for the MEG dataset and two for the fMRI dataset). Two of these initial estimations, used for the EVC-fMRI data and the early-MEG window, were based on purely perceptual features (i.e. edges) while the other two, used for the ITC-fMRI data and the late-MEG window, were based on the categorical information extracted from the training data. The RDMs predicted by these initial models were then improved by weighted average using the RDMs derived from the DNNs features that were provided (see below).

## Initial model for perceptual RDMs

The initial model for the perceptual RDM was based on three steps. First, images from the training and test sets were resampled to have the same number of pixels. Second, edges were extracted and finally gaussian smoothing was applied. Figure 1 shows the result of these transformations in one of the images of the training data.

**Figure 1.**
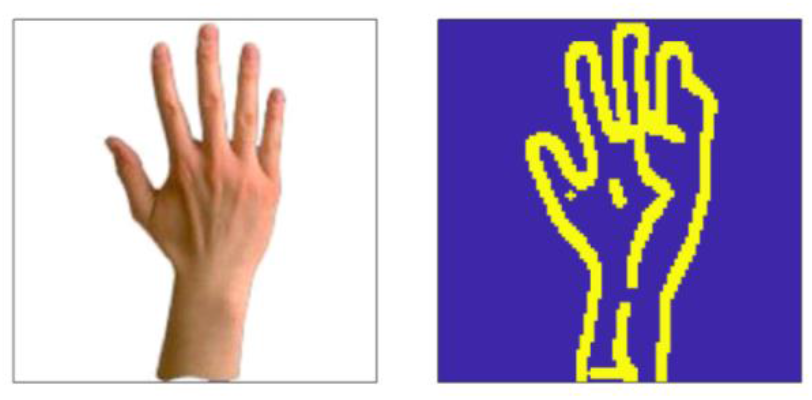
The figure shows the features extracted for one image in the 92 training set.

A (perceptual) RDM for a given set of images was obtained using as distance metric the overlap between the images after the feature extraction process (i.e. one minus the sum of the number of pixels overlapping between any pair of images).

The number of pixels (*n*) used for the resampling procedure, the threshold for image edge extraction (*t*) and the size of the gaussian smoothing kernel (*s*) were optimized by maximizing the correlation between the predicted RDMs and the provided RDMs from the training data set. The resulting parameters were *n* = (100 × 100) pixels, *t* = 12 and *s* = 1 for the early-MEG window data and *n* = (166 × 166) pixels, *t* = 13 and *s* = 2 for the EVC-fMRI data.

## Initial model for categorical RDMs

We manually labelled the images in the training data set with 92 images as belonging to eight categories (note that only the training set was used): 1) objects-scenes, 2) animals, 3) human, 4) fruits-veggies, 5) faces, 6) hands, 7) monkeyfaces, 8) animal-faces. We then trained a gaussian naïve bayes (GNB) classifier to distinguish between these categories on the basis of the representation of the images by the fully connected layer of the vgg-FC8 network (1000 features). Prediction accuracy reached 90.22% using leave one-out cross validation in the 92-images set (Figure 2).

**Figure 2.**
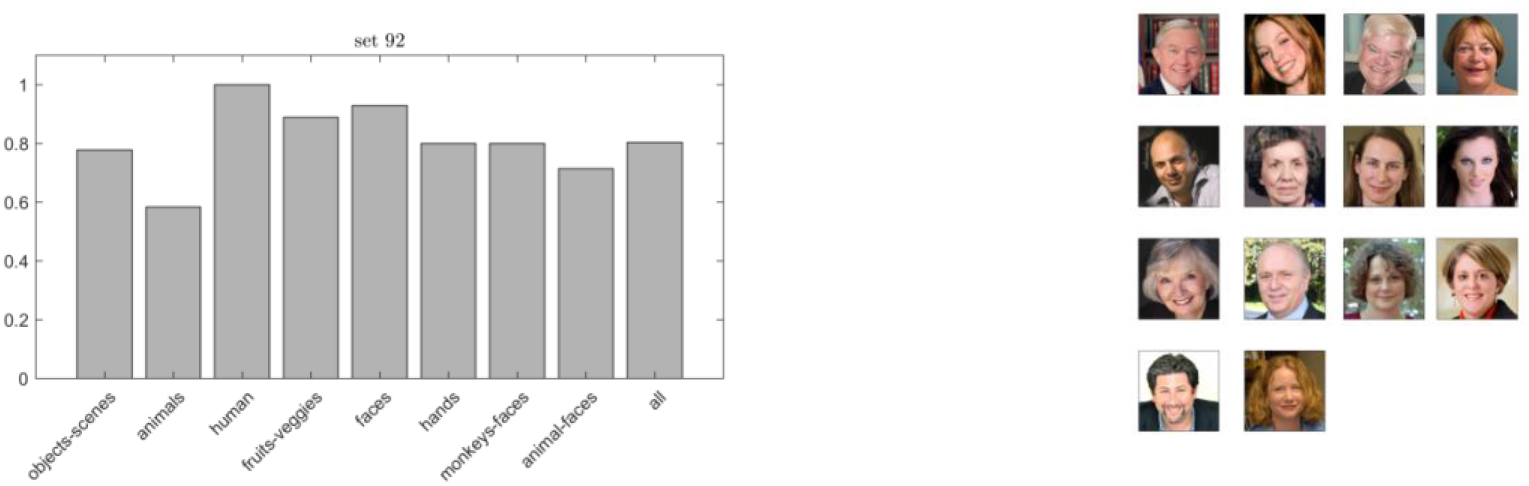
(Left) Average leave-one-out classification performance on the 92-image set (per category and in average across categories). (Right) images in the test data that are predicted to be faces.

The 92-image set was then used to obtain a “categorical” (8 × 8) RDM by averaging the values of the initial 92 × 92 RDMs within blocks of images belonging to the same category. This categorical-RDM was different for the late MEG data and the ITC fMRI data.

The test set images were assigned to one of the eight categories using the trained GNB classifier. Inspecting the result showed a high consistency among the test images classified as belonging to the same class (see e.g. on the right of Figure 2 we display the test set images that are predicted to be faces). To predict the RDM for the test set, we defined the distance between any pair of images to be equal to the distance between the corresponding predicted categories in the 8 × 8 RDM derived from 92 image training set. Note that the categorical structure of the test data was only inferred by the classification algorithm based on the training data and not imposed a-priori.

## Improvement in model representations using DNN features

The contribution of DNNs was included in the final perceptual and categorical models via a weighted average between the initial RDM (*R*^0^) and an RDM derived from the DNNs (*R*^*DNN*^):

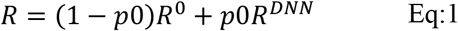

We considered each of the provided DNNs separately. The *R*^*DNN*^ matrices for the different networks were obtained for every block by averaging the activations across all features (e.g. conv1 in alexnet contains 64 feature maps of size: 55, 55). Next the RDMs were obtained using the spearman correlation coefficient between the averaged maps for every pair of images. The weighting contribution (i.e. the parameter *p*0 in Eq. 1) was optimized in the training set for every modality and region (MEG-early MEG-late, fMRI-EVC and fMRI-ITC) separately. Figure 3 shows for each of the three networks provided the performance relative to the noise ceiling of every block as the contribution varies from *p*0 = 0 to *p*0 = 1 for the fMRI-EVC in the 92 (upper) and 118 (bottom) images set. Selecting those blocks that provided the largest improve in the training data the initial RDMs for the test data were combined with those RDMs derived from the DNN for the test set images.

**Figure 3.**
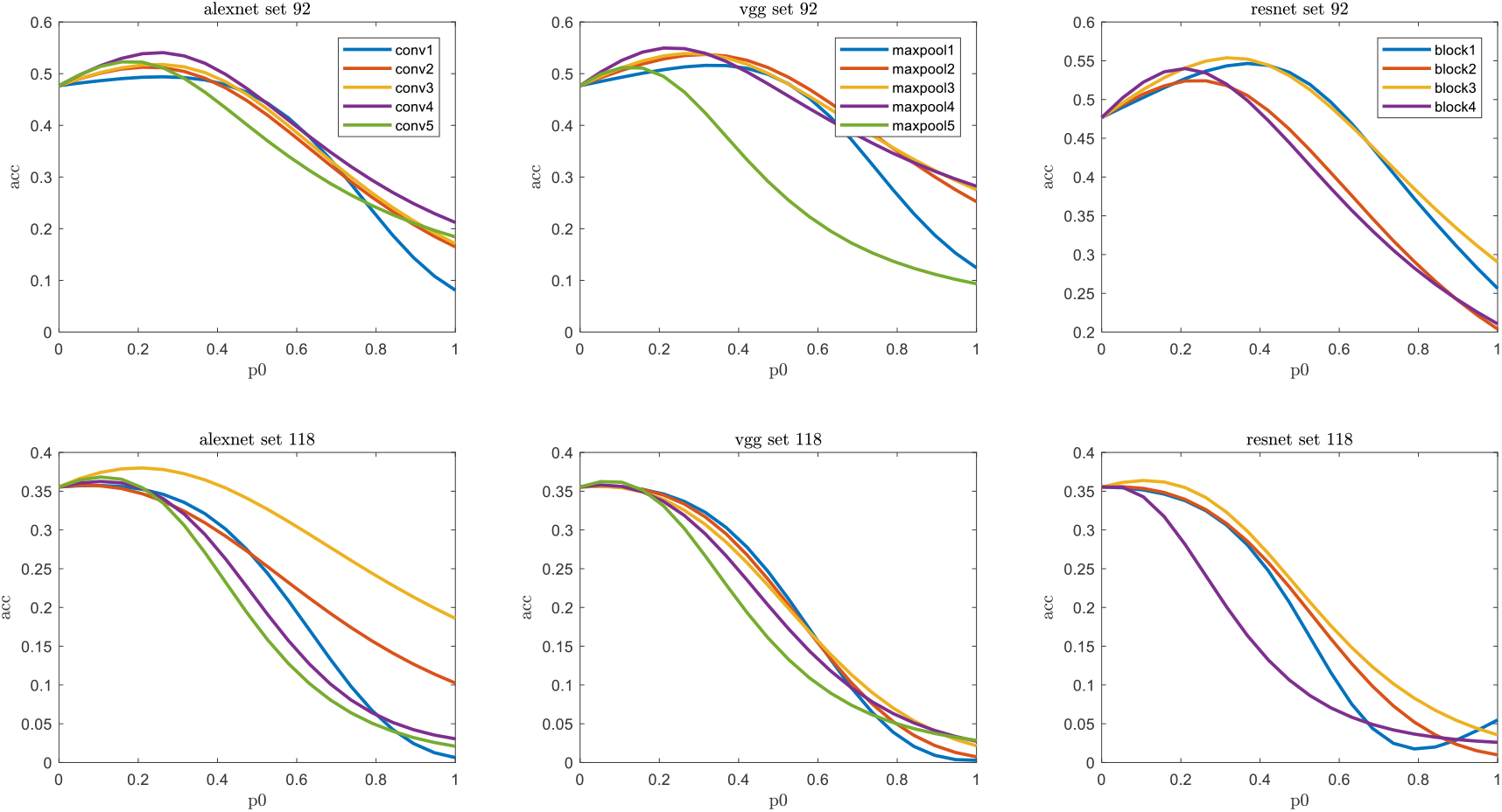
The figure shows the model performance relative to the noise ceiling for an increasing proportion of the DNN contribution into the RDMs (see Eq.1). This analysis corresponds to fMRI-EVC dataset using the 92 and the 118 data sets.

## Results and Discussion

Figure 4 (top row) shows the predicted RDMs for the initial perceptual (used for MEG-early and fMRI-EVC) and the categorical (used for MEG-late and fMRI-ITC) RDM estimations. The adjustment obtained by the weighted averaged of these initial representations with the DNNs based RDMs is visualized in the bottom row of Figure 4.

**Figure 4.**
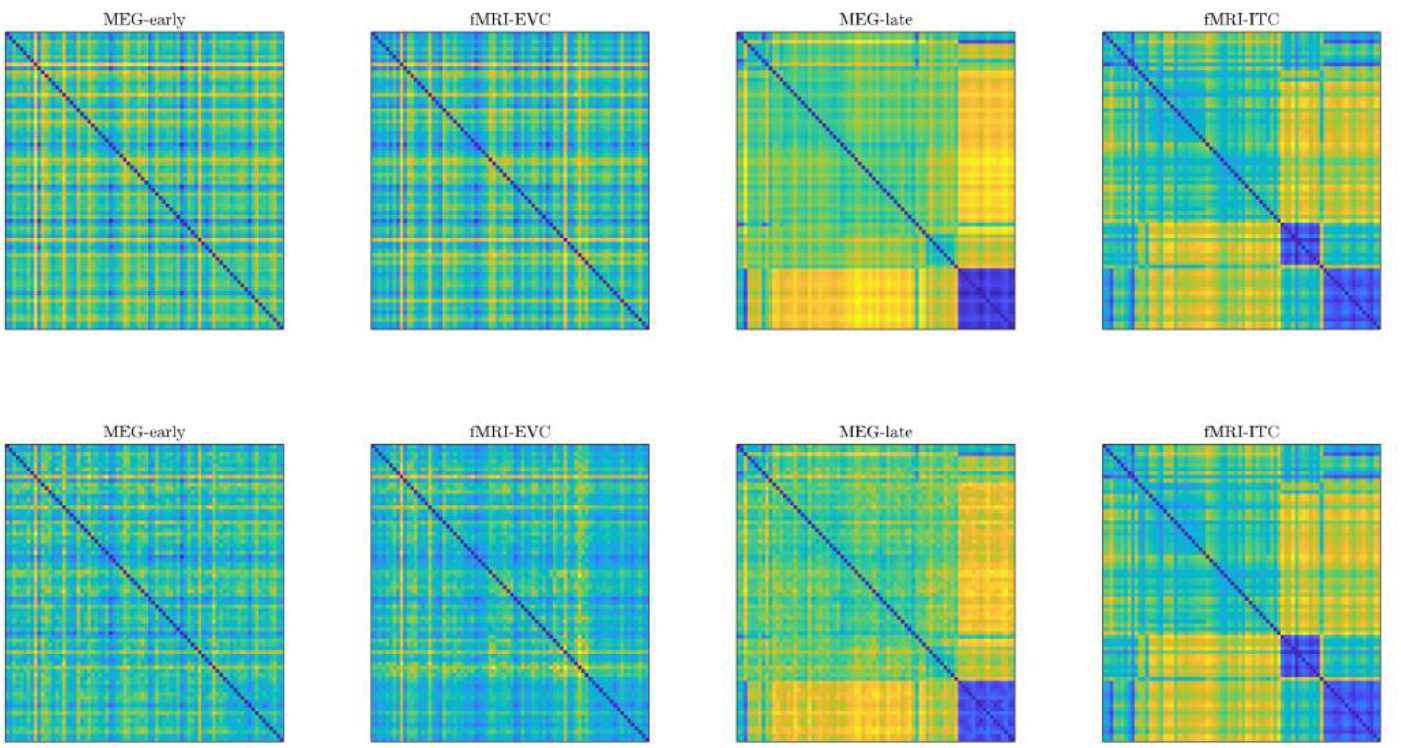
Top row: Initial RDMs estimation for MEG-early, fMRI-EVC, MEG-late and fMRI-ITC. Bottom row: corresponding RDMs obtained after including the contributions of the DNNs.

In Table 1 we present, for both modalities and regions of interest, the noise normalized explained variance for the test data using the initial RDM estimation and the improvement obtained when the information from the DNN was added to the prediction. The reported values are based on submitting our predictions to the online system.

**Table 1:** Contributions of the initial RDMs and the DNN-RDMs to the model performance.

| | fMRI Noise Normalized $R.^2$ (%) | | meg Noise Normalized $R.^2$ (%) | |
| --- | --- | --- | --- | --- |
|  | EVC | ITC | Early | Late |
| Perceptual + Categorical | 25.20 | 20.99 | 45.28 | 44.43 |
| Perceptual + Categorical + DNN | 32.88<br>vgg-L3<br>vgg-L4<br>vgg-L5 | - | 50.95<br>alexnet<br>L3,L5 | 53.59<br>alexnet L5<br>vgg L5<br>resnet L4 |

The initial RDM estimation already captures a large proportion of the total noise normalized explained variance that was achieved by the combined model (that includes DNNs). Except for ITC all the RDM predictions improved with the contributions from DNNs. However, the improvement obtained from the DNNs observed in the training data was not often reproduced in the test data when the predictions were evaluated via the online submission system. Among the provided DNNs, the vgg network was the one that contributed the most to improve the predicted scores.

